# Enhanced Sampling Simulations of RNA-peptide Binding using Deep Learning Collective Variables

**DOI:** 10.1101/2024.08.01.606277

**Authors:** Nisha Kumari, Sonam Dhull, Tarak Karmakar

## Abstract

Enhanced sampling (ES) simulations of biomolecular recognition such as binding of small molecules to proteins and nucleic acids targets, protein-protein association, and protein-nucleic acids interactions have been gaining significant attention in the simulation community due to their ability to sample long timescale processes. However, a key challenge in implementing collective variable (CV)-based enhanced sampling methods is the selection of appropriate CVs that can distinguish the system’s metastable states and, when biased, can effectively sample these states. This challenge is particularly acute when simulating the binding of a flexible molecule to a conformationally rich host molecule, such as the binding of a peptide to an RNA. In such cases, a large number of CVs are required to capture the conformations of both the host and the guest, as well as the binding process. In our work, we employed the recently developed Deep Targeted Discrimination Analysis (DeepTDA) method to design CVs for the study of the binding of a cyclic peptide, L22 to a TAR RNA of HIV as a prototypical system. These CVs were used in the on-the-fly probability-based enhanced sampling and well-tempered metadynamics simulations to sample reversible binding and unbinding of L22 peptide to the TAR RNA target. The enhanced sampling simulations revealed multiple binding and unbinding events, which enabled the calculation of the free energy surface for the peptide binding process. Our results demonstrate the potential of the DeepTDA method for designing CVs to study complex biomolecular recognition processes.

## INTRODUCTION

Sampling of rare events, such as small molecule binding to proteins and nucleic acid targets, protein-protein association, protein-nucleic acid interactions, and conformational changes in biomolecules, is one of the long-standing problems in structural biology and biophysics. Molecular dynamics (MD) simulations, skilled to simulate a system for only a few milliseconds present a great challenge to learn the insights into atomistic detail behind rare events that occur beyond a few milliseconds timescales. The ability to sample a greater area of the configuration space of complicated systems within affordable simulation time and to increase the utility of MD simulations have been made possible by enhanced sampling (ES) techniques. By assisting in removing significant energy barriers and allowing the exploration of system states that are normally unattainable in brute-force MD simulations, these ES methods also help calculate the thermodynamics and kinetics of a physicochemical and biological process. Over the years, numerous ES methods have been developed to address the MD simulation timescale issue. These methods can be chiefly divided into three categories: Biasing approaches,^1–3^ adaptive sampling methods,^4^ and generalized ensemble methods.^5^ The methods that involve supplementing the system’s Hamiltonian with an external bias potential which is a function of the atomic coordinates, among the many other alternatives, have been exhaustively used by the simulation community.^1,6,7^ In all practical purposes, in the latter category, the external bias is defined as a function of the system’s slow degrees of freedom, often termed as collective variables (CVs). The bias enhances the CVs fluctuations and thereby removes kinetic bottlenecks by favoring transitions between different metastable states.

CVs have traditionally been constructed out of physical-chemical intuition. For instance, in the case of an RNA-ligand system, the center of mass distance between RNA and the ligand can be chosen as a CV to sample the ligand binding and unbinding processes. However, it is possible to miss significant slow degrees of freedom that might impede convergence when one moves forward in this manner. Additionally, the intricacy of the problems that can now be simulated necessitates an alternative, more automated methodology in which variables are taken straight out of the MD simulation data.^8^ Selecting CVs based on data may be seen as a manifold learning issue, where the objective is to choose a low-dimensional spectrum that best captures the key slow dynamics and or dominating metastable states of the system. Machine learning (ML) techniques are excellent at inferring patterns and intricate functions straight from data, and this is the ideal setting for them to work. A variety of physical, chemical, and biological processes have been studied and accelerated *via* the use of data-driven collective variables, proving the significant value these methods may have for atomistic simulations.^9–26^ The recently developed Deep Targeted Discriminant Analysis (Deep-TDA)^27^ has shown to be effective among many of these techniques. The CVs as functions of numerous physical descriptors have recently been learned by this technique. ^28^ Data from short unbiased simulations carried out in the metastable basins is used to construct the DeepTDA CV. This method is quite appealing since the number of descriptors that may be employed is theoretically unlimited, and the CV generation process is semi-automatic.

In our work, we employ the Deep-TDA method to develop effective CVs for the study of a cyclic peptide (L22) binding to the HIV TAR RNA (see Figure 1). The L22-TAR RNA is an interesting system since both the host (RNA) and the guest (L22) are fluxional, and it is a non-trivial task to design a suitable CV to sample the binding/unbinding events using ES simulations. The interaction between the HIV-1 trans-activator Tat protein and its associated trans-activation response element (TAR) RNA is an essential component of viral transcription and replication.^29,30^ It has been demonstrated that interfering with Tat’s binding to TAR RNA is a particularly appealing tactic for preventing HIV infection. While nuclear magnetic resonance (NMR) and X-ray investigations can provide precise descriptions of the structures of both the bound and free states of RNA and protein, the mechanism of binding is usually not well understood. ^31^ Significant conformational changes in the molecular system are often caused by the interaction of RNA and protein.^32–34^ RNA-protein interactions are highly influenced by conformational flexibility, even though its biological role is yet unclear. The difficulty in designing RNA-binding drugs stems from our limited understanding of RNA/ligand molecular recognition events.^35–38^ These events are primarily driven by dynamically induced fit mechanisms, which are defined as ligand-induced conformational rearrangements of local RNA secondary structure elements that stabilize a specific conformation of the RNA-protein complex.^34,39^ The understanding that an effective CV should be able to discern between the starting and final states forms the foundation of our work. Our primary focus in this work is to calculate the free energy difference (ΔG) between the bound and unbound states, as well as the primary residues that are crucial for the binding and recognition of the L22 cyclic peptide to the TAR RNA groove. Thus, we employ the recently developed Deep-TDA approach to obtain a CV that can effectively capture the binding and unbinding event of the cyclic peptide.

**Figure 1:**
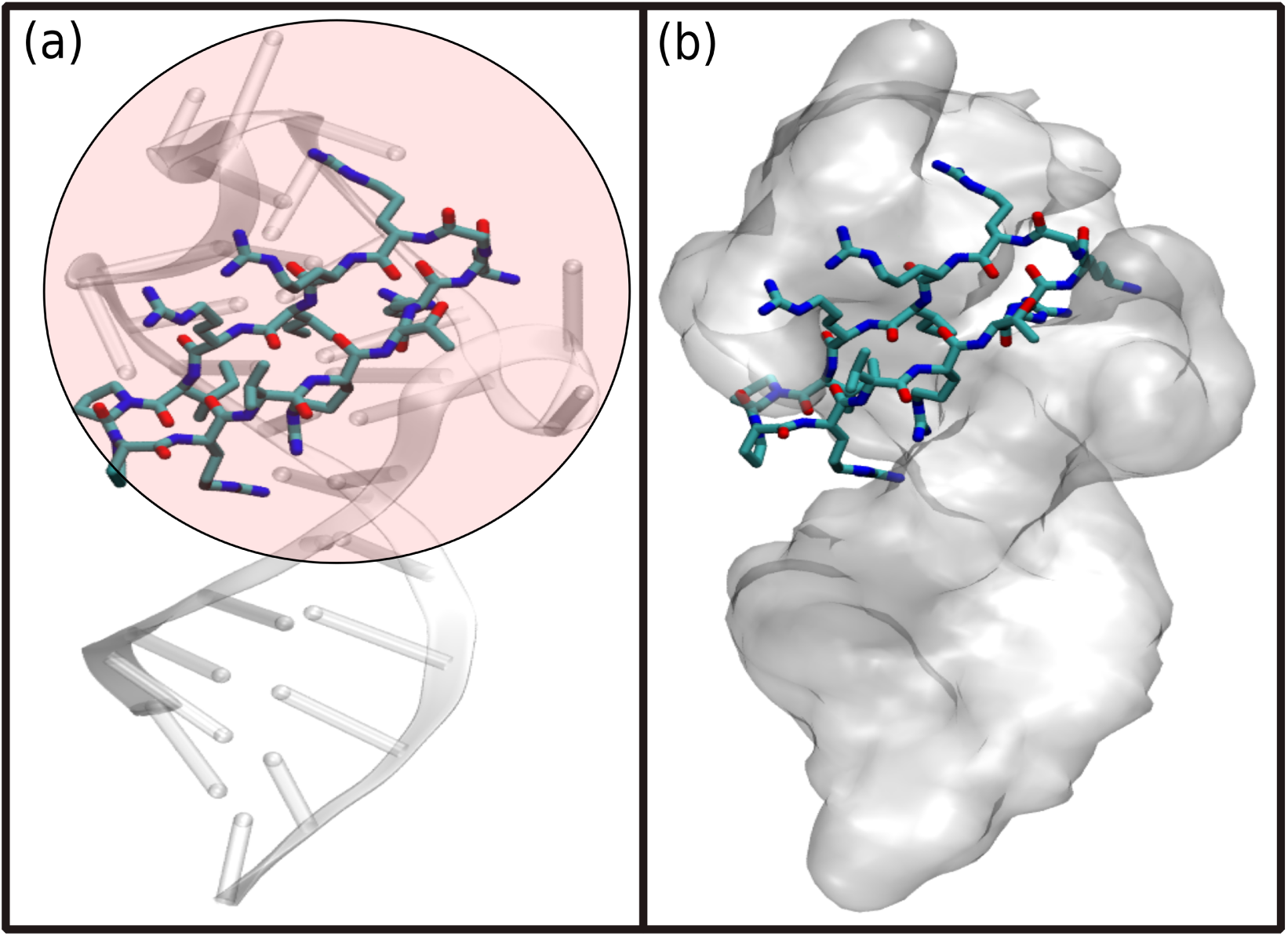
HIV TAR RNA in complex with L22(cyclic peptide mimic of TAT protein). The L22(represented in licorice format) is bound to the apical loop of the TAR RNA. The TAR RNA is represented in NewCartoon in (a) and solid Surface in (b). The apical loop region of the RNA is shaded in pink. The region down the apical portion is the base pairing region.

## COMPUTATIONAL METHODS

### Enhanced sampling method

As slow convergence rates for complicated systems limit the range of metadynamics simulations, achieving multiple binding-unbinding events for complex systems is still a tedious job. Here we employ the recently developed On-the-fly Probability Enhanced Sampling (OPES) method to study the binding/unbinding of the peptide to/from the TAR RNA. In OPES simulations, a Gaussian bias is periodically deposited to enhance the fluctuations of the CV space. For complex systems such as our case where the host and the guest both show large conformational flexibility, choosing the right set of CVs is a challenging task. We choose CVs based on the available crystal structure and chemical/physical intuitions. However, when it comes to dealing with a complex biomolecular system, a large number of CVs are required to describe the system’s metastable states in our case, peptide bound to the RNA and the unbound structure. Dealing with a large number of variables is not practical for running ES simulations. One could in principle combine them linearly and obtain a CV. 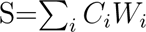, where *C_i_* are the contacts and *W_i_*are the weights. Our initial trial with this CV along with the RMSD of RNA, indeed resulted in a trajectory in which we saw one unbinding event. However, soon we realized that the linear combination is not effective in our case. Then we switched to combining the contacts non-linearly, and for this purpose, we used the Deep-TDA method.^27^

### Deep-TDA

The Deep-TDA approach was developed to facilitate data-driven CV design. It commences with the description of a system’s metastable states. Using a wide range of physical descriptors as inputs gathered through short unbiased simulations that solely sample the metastable basins. The Deep-TDA CV is built on a discrimination criterion of Fisher type, and the CV is expressed directly as an output of a feed-forward neural network (NN). When dealing with a two-state situation, the NN is trained to ensure that the metastable state distribution along the CV follows a preassigned bimodal Gaussian distribution. One of the two Gaussians describes the distribution of the A configurations, while the other Gaussian describes the B configurations (Figure 2(b)). The target Gaussian distributions have preassigned positions and widths. A Fisher-like ratio (equation 1) can be used to gauge how well a target distribution can distinguish between the two states.

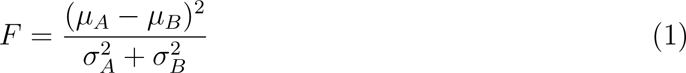

**Figure 2:**
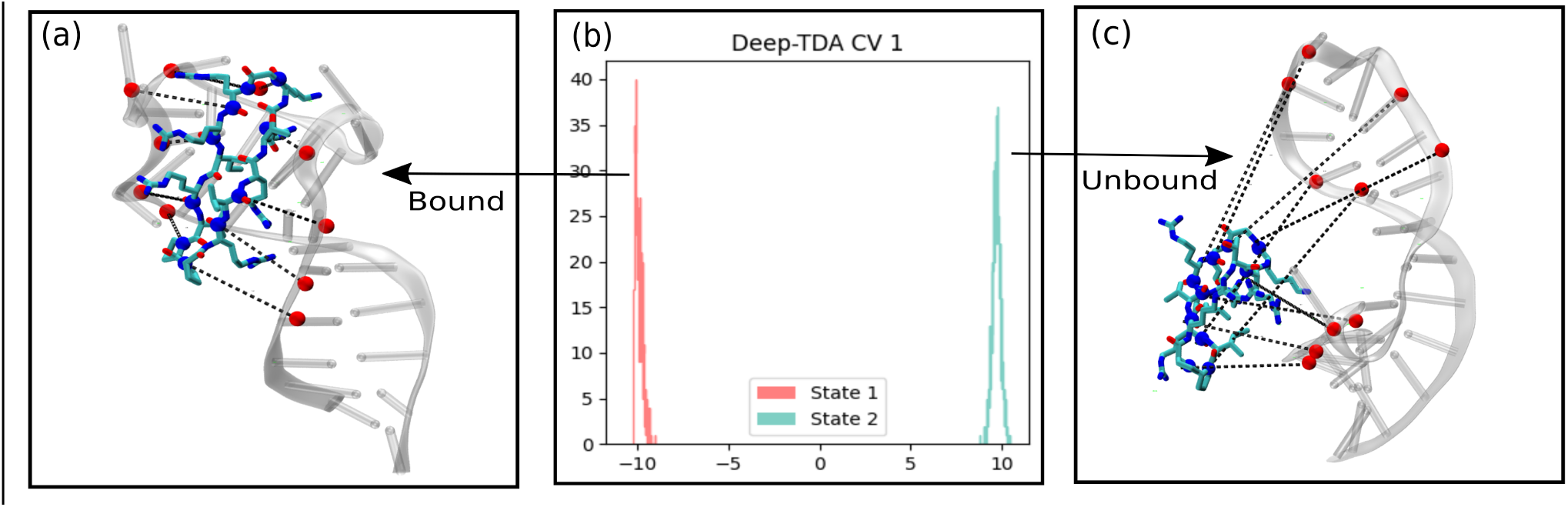
(a) Contact pair distances as input descriptors from the bound state configuration, (b) bimodal Gaussian distribution was achieved after training physical descriptors as inputs gathered through a short unbiased simulation from both the bound and the unbound state, and (c) Contact pair distances as input descriptors from the unbound state configuration.

where *µ_A_* and *µ_B_*are the average values and 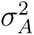 and 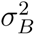 represent the variances for the two states in the CV space. However, it is appropriate to employ 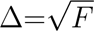 as a parameter to characterize the objective function, since F grows rapidly to very large values when the spacing between the states is raised. The details of the DeepTDA method can be found in Ref. 27

We employed the OPES method (explore variant) to sample binding and unbinding events and calculate the binding free energy surface (FES). The Deep-TDA CV (S1) along with a second CV (S2), which is the root mean square displacement (RMSD) of the apical-loop region of the TAR RNA was used in the OPES simulations to explore the conformational space of the system. To train the deep-TDA CV in both the bound and unbound states, a number of contact pair distances between the cyclic peptide and TAR RNA were selected based on observation of equilibrium simulation physical intuition (Figure 2a, c). Then, leveraging these descriptors to build the Deep-TDA CV, an unbiased simulation was run in both the bound and unbound states to produce the training data. The Pytorch ^40^ library was used to train the model and further, the trained model from Pytorch was imported to PLUMED [using the interface developed by Bonati et al.^9^ The GROMACS-2021.4 MD engine^41^ patched with PLUMED2.9,^42^ was used to run the ES simulations.

## RESULTS

Equilibrium simulations of the free RNA, free cyclic peptide, and RNA-cyclic peptide complex systems in water were carried out for 2-microsecond (*µ*s) each. The conformational dynamics of RNA and peptide both in complex and in the free states were calculated using root mean square displacement (RMSD) (Figure S1 in the Supporting information (SI)). The RNA is found to be conformationally flexible in its free form with an average RMSD fluctuating below 5Å. A reduction in the conformational flexibility was observed in the TAR RNA-peptide complex state with an average RMSD fluctuating around 4Å. The nucleotides spanning the base pairing area varied the least while the apical loop region (Figure 1b) of the RNA, which served as the RNA’s primary binding motif, contributed most to the overall RNA fluctuations. The L22 cyclic peptide, on the other hand, was highly stable in its free state with an average RMSD below 1Å. For the cyclic peptide in complex with RNA, a slight conformational shift in the RMSD of the peptide was observed after 1.1 *µ*s. The peptide slid down away from the binding motif toward the base pairing region, and the conformation of the peptide tilted slightly upright, as was observed from the simulation trajectory.

The conformational shift induced in the cyclic peptide can be attributed to the flexibility of the apical loop region of the RNA. Despite the conformational shift and the movement of the peptide slightly away from the binding site, it was found to still strongly intact with the RNA due to the numerous electrostatic interactions present between the RNA and the peptide. To find out the fluctuations of the peptide at the residue level, root mean square fluctuation (RMSF) for all the residues of the cyclic peptide was calculated (Figure S2 in SI). The ARG residues flanked on the peptide were majorly involved in the interaction with the RNA backbone and, hence had the highest fluctuations. As can be seen from the RMSF plot, residues comprising VAL, THR, GLY, ILE, and PRO have comparatively lesser fluctuations. Overall, the fluctuations of the RNA was reduced in the complex state, and the cyclic peptide conformationally reoriented itself to fit in the binding groove of the RNA.

On visualization of the simulation trajectory, the residue-level interactions between the RNA and the peptide that led to the complex’s stability were observed (Figure 3). The conformational shift of the peptide after 1.1 *µ*s resulted in the overall shift of the interactions between the peptide and the RNA. The RNA backbone phosphate groups form crucial salt bridges with the cyclic peptide ARG residues, and these interactions (ARG1-G21, ARG3G37, and ARG11-A22) were found stable throughout the entire simulation time. The cyclic peptide’s reorientation in the binding groove causes the ARG residues to interact with other nucleotides, and they frequently exhibit several salt bridge formations at once. Additionally, the side chains of ARG5, ARG3, and LYS6 residues form hydrogen bonds with nucleobases G28, G26, and C30, respectively. A continuous shift between hydrogen bond to salt bridge formation was observed for the LYS residue. Apart from hydrogen bonding, ARG residues were found to interact with nucleobases of G34, A22, and U23 *via* van der Waals and cation*π* (on geometric basis) interactions thereby exerting additional stabilization to the peptide bound to TAR RNA. A few representative interactions are shown in Figure 3.

**Figure 3:**
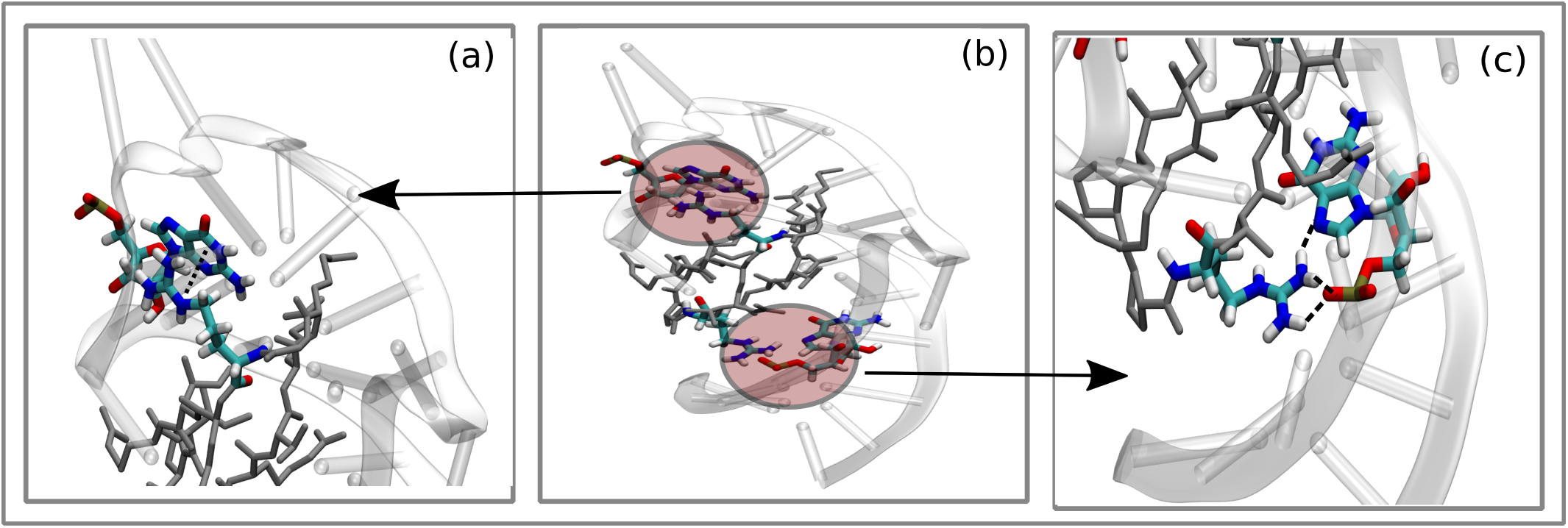
(a) *π*-cationic interaction between ARG8 residue of cyclic peptide and the G34 nucleobase of RNA, (b) Configuration profile of the complex incorporating various kinds of interaction in a single frame, and (c)electrostatic interaction as well as hydrogen bonding between peptide ARG1 residue and G21 nucleotide of the RNA

From long microseconds equilibrium simulations, the key interactions stabilizing the peptide inside the RNA group have been realized. To determine if the cyclic peptide binds spontaneously to the RNA’s peptide binding site, we carried out a 10 *µ* unbiased MD simulation in which the cyclic peptide was initially kept approximately 40Å away from the RNA’s binding groove. The cyclic peptide approached the apical loop region directly and got attached to the top of the apical loop portion of the backbone in an upright fashion. ARG residues flanking the peptide helped it get partially attached to the RNA. The peptide stayed there in the semi-bound state for over 5-microsecond simulation time. Even though it did not leave the partially bound site, it did not penetrate any further into the actual binding site (crystal structure). To quantify the extent of binding, we computed the linear sum of multiple contact pairs between RNA and peptide atoms and compared the results with those obtained from the equilibrium simulations of the complex state (Figure 4). As evident from Figure 4, not all the connections were built during the binding process which indicates that the cyclic peptide remained partially bound throughout the simulation. We anticipate there is high free energy barrier that prevents the cyclic peptide from penetrating the groove and reach to its actual bound state.

**Figure 4:**
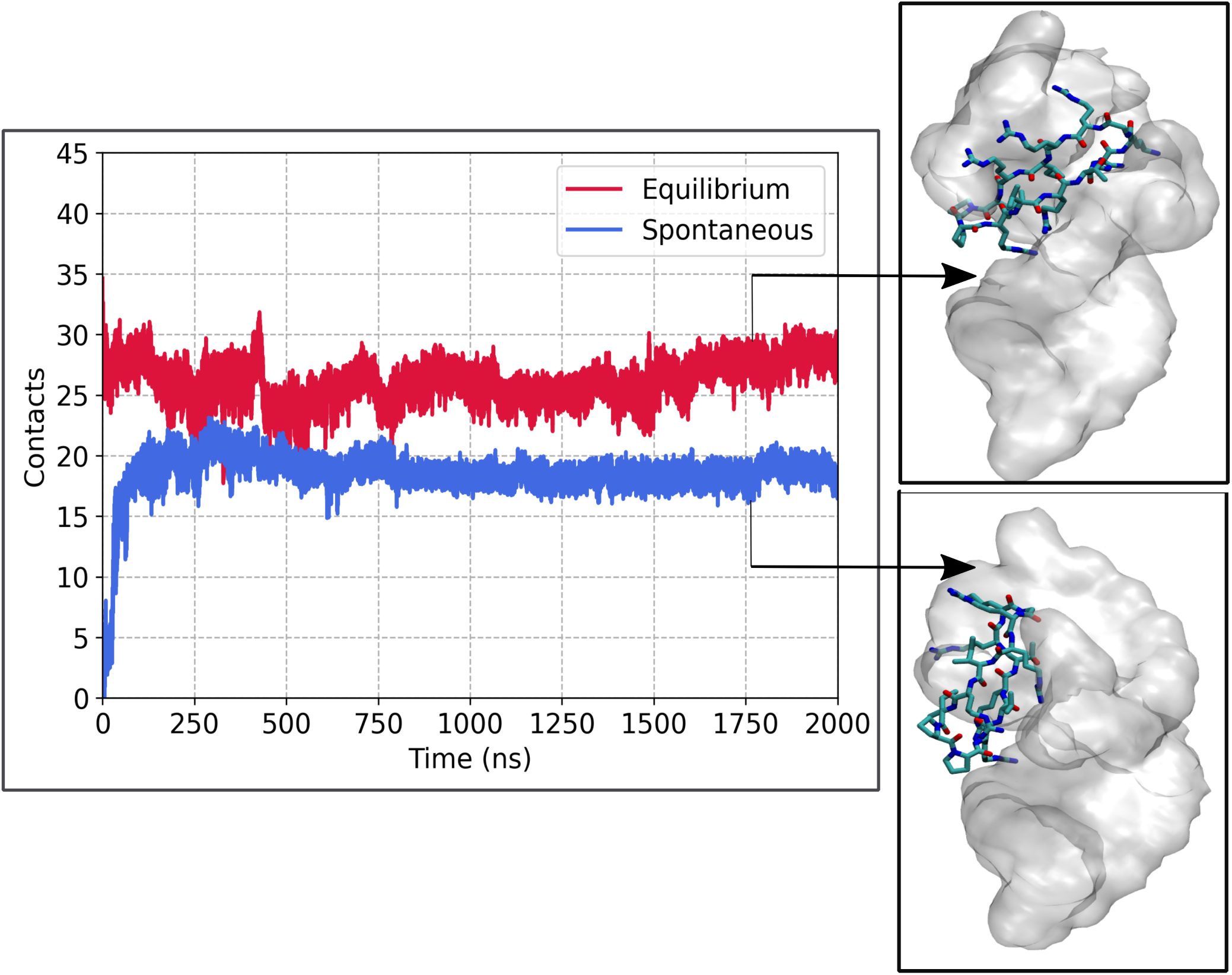
The graph plots the spontaneous binding of the cyclic peptide to the RNA vs the equilibrium trajectory over 2(*µ*s) simulation run. The y-axis incorporates contact pair distances between the RNA apical loop region and the cyclic peptide. The blue plot corresponds to the spontaneous trajectory while the red plot for the equilibrium trajectory. Their representative figures correspond to the configuration at 2(*µ*s) simulation time

To fully comprehend peptide-RNA binding/unbinding events, we switched to OPES simulations. We have used two CVs the first one is the Deep-TDA which is a non-linear combination of forty-one contact pair distances, mostly consisting of distinct combinations of electrostatic interactions, hydrogen bonds, and long-range contacts between the RNA and the peptide (Figure 2), and the second CV is the RMSD of the RNA apical loop shaded in pink (Figure 1). Multiple binding-unbinding transitions were observed throughout the 5 *µ*s OPES simulation (Figure S3 in SI), which demonstrates an efficient sampling of the metastable states. In the CV profile S3, -10 in the y1-axis is the initial bound state, and 10 is the unbound state, while states between -10 to 10 are other conformations of the complex. Conversely, the y2-axis represents the RNA apical loop RMSD variations as the system explores various metastable states. We started our simulation from the bound state where all the 41 contact pair distances were intact. The cyclic peptide loses all of its connections and starts to drift away from the binding site within a few tens of nanoseconds. Following its detachment from the RNA, the cyclic peptide gradually returns to the binding groove where, in the partially bound state, it reorients itself to bind in its initial configuration (Figure 5a-f). During binding, the peptide ARG residues function as recognition motifs, drawing the peptide to the binding location by strong electrostatic interactions.

**Figure 5:**
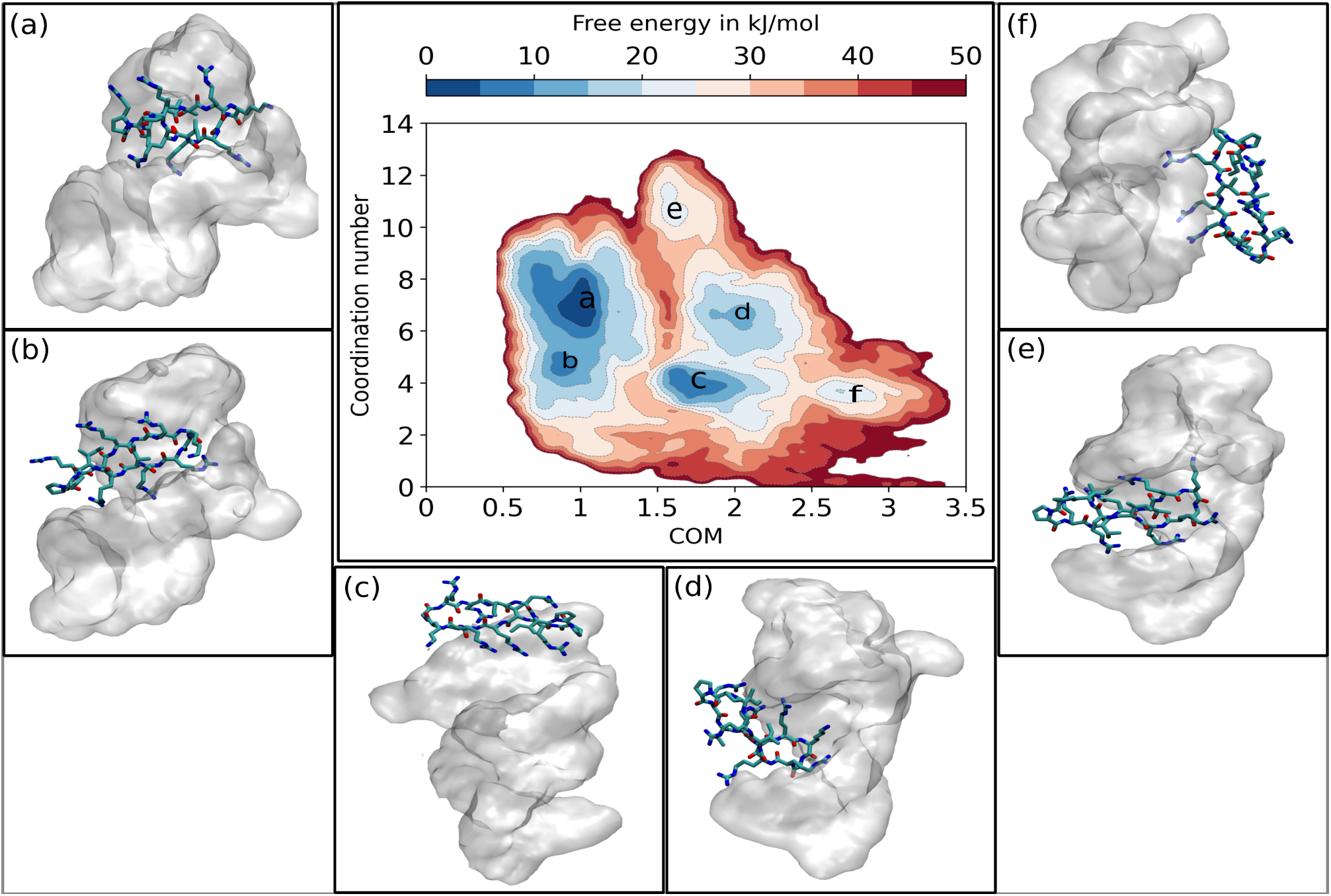
Free energy landscape projected along a number of phosphates in the RNA backbone that coordinate with the positively charged nitrogen atoms in the arginine and lysine of the cyclic peptide and center of the mass distance between cyclic peptide and the RNA apical loop region. Basin (a) corresponds to the initial bound state that is being visited twice during the whole simulation time, while other basins represent the different stable conformational states exhibited by the complex during OPES simulation.

The bias deposited to enhance the fluctuations of the CV was reweighted to construct the FES as a function of CVs. Since a fair amount of contact pair distances chosen comprised electrostatic and H-bond distances between RNA phosphate and peptide positively charged ARG atoms, any motion in the side chains of ARG residues led to a shift in the FES profile. Thus, we used two physical descriptors to project the free energy and assign the bound and unbound states of the peptide (Figure 5a-f). The first is the separation between the centers of the cyclic peptide and the RNA apical loop region, and the second is the number of phosphates in the RNA backbone that coordinate with the positively charged nitrogen atoms in the ARG and LYS of the cyclic peptide (Figure 5). The free energy landscape spanning the entire conformational space traversed during the 5(*µ*s) OPES simulation run primarily comprises six major basins represented as a, b, c, d, e, and f. (Figure 5) The basin a having the lowest FE resembles the peptide bound RNA crystallographic structure. This conformational state was visited twice after the peptide fully unbound from the RNA as is evident from the CV profile in Figure S3. Of the other minimum energy basins, state b corresponds to the bound state with slightly higher RNA apical loop RMSD, as when the apical loop of RNA opens up, the cyclic peptide drifts a little away from the binding groove resulting in losing some of the phosphate coordination. The basin c is a partially bound state close to the binding groove where the cyclic peptide is attached to the upper backbone of the apical portion of the RNA, and this state is separated from the bound state by an energy barrier of *∼*30 kJ/mol. Basin d and e are other binding poses acquired by the complex when the peptide binds to the RNA base pairing region, also basin e is the state where the phosphate coordination is reached to maximum. Even though the number of electrostatic interactions is higher in this region, free energy is higher in basin a. In other words, even though the phosphate coordination is high in basin e, basin a is the minimum energy state indicating the importance of the apical loop region. The RNA and cyclic peptide when bound in the apical loop region is conformationally less flexible exhibiting a lower entropy and a higher stability in comparison to the state when it is bound to the base pairing region where a binding groove is missing and hence is prone to high flexibility and hence less stability. Finally, the peptide attaches to the rear side of the base pairing area down to the apical loop binding groove, resulting in basin f. Further, the high selectivity of the crystallographic state can be attributed to the binding pose, other significant interactions discussed earlier, and the constrained conformational flexibility of the peptide which minimizes the entropic penalty during binding. Thus, given the conformational space, a significant number of states were explored incorporating deep-TDA and OPES simulation methods at a significantly lesser time and computational cost.

## CONCLUSION

In summary, our study focuses on the use of DeepTDA-based collective variables in On-the-Fly Probability Enhanced Sampling (OPES) to simulate multiple binding-unbinding events for a complex peptide-RNA system. The relative free energy was calculated which allowed us to locate all the major basins representing different conformational states of the peptide-RNA complex. The free energy landscape calculated from the OPES simulations revealed the lowest free energy corresponds to the peptide-bound NMR crystallographic state. De-signing CVs for fluxional systems is an inherently challenging task, and our results indicate deep learning methods to be valuable for understanding complex biomolecular processes. However, despite the success of the current approach, the intricate biomolecular systems necessitate further refinement and enhancement of these deep-learning methods. Further research should focus on optimizing these deep-learning techniques to achieve greater accuracy and robustness in CV design and deepen our understanding of biomolecular interaction and transition.

## Supporting Information Available

The supporting information contains

RMSD plot(peptide in water, RNA in water, peptide in complex with RNA, RNA in complex with peptide); RMSF of cyclic peptide; deep-TDA CV plot.

## TOC Graphic

## Notes

### Competing Interest Statement

The authors have declared no competing interest.

