## Supplemental Information for "Enhanced Sampling Simulations of RNA-peptide Binding using Deep Learning Collective Variables"

### RMSD and RMSF

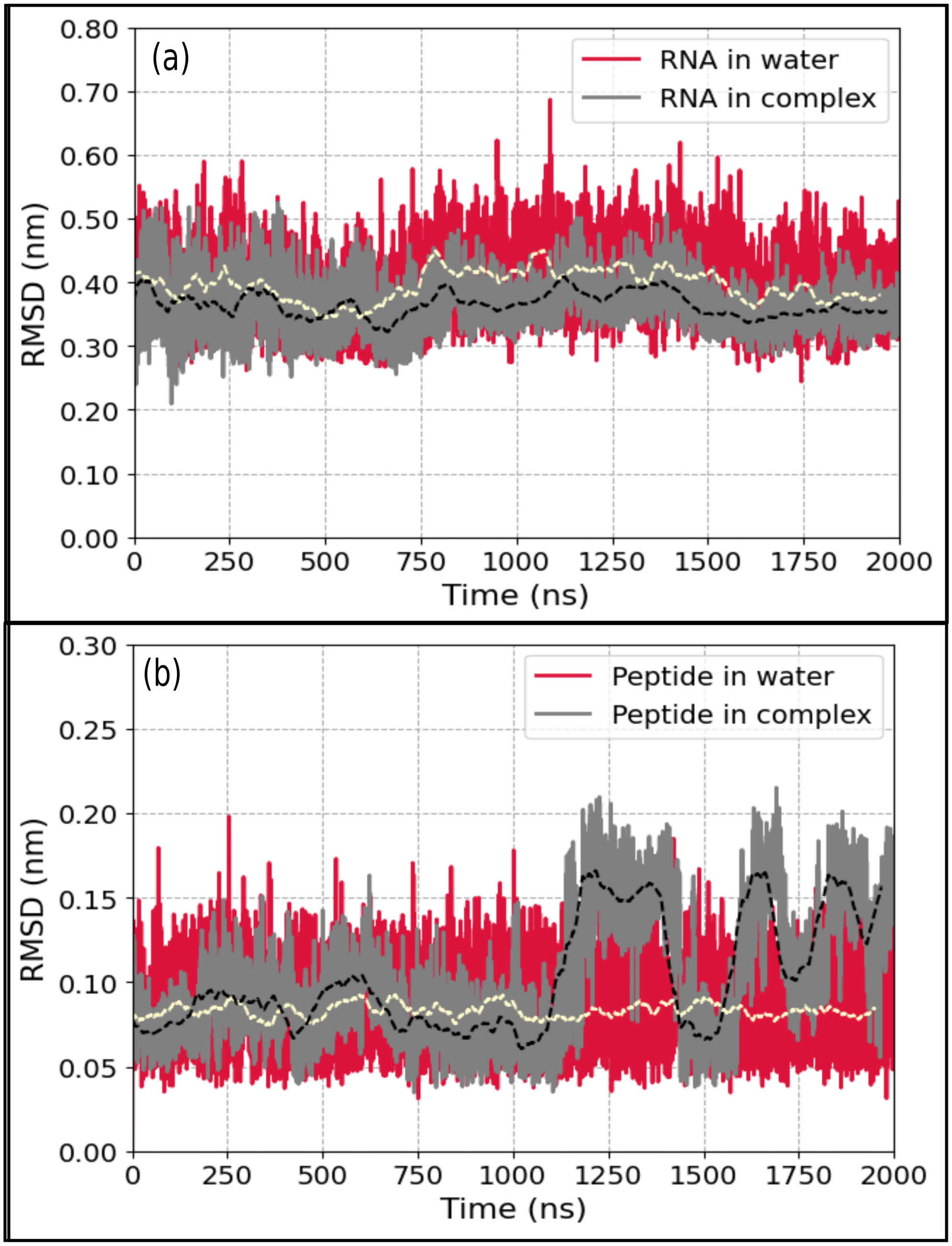


Figure S1: (a)RMSD plot of RNA in water(crimson) and RNA in complex with cyclic peptide(grey). (b) RMSD plot of peptide in water(crimson) and peptide bound to RNA(grey). The running average for RNA and peptide in water is shown in yellow while for complex it is shown in black.

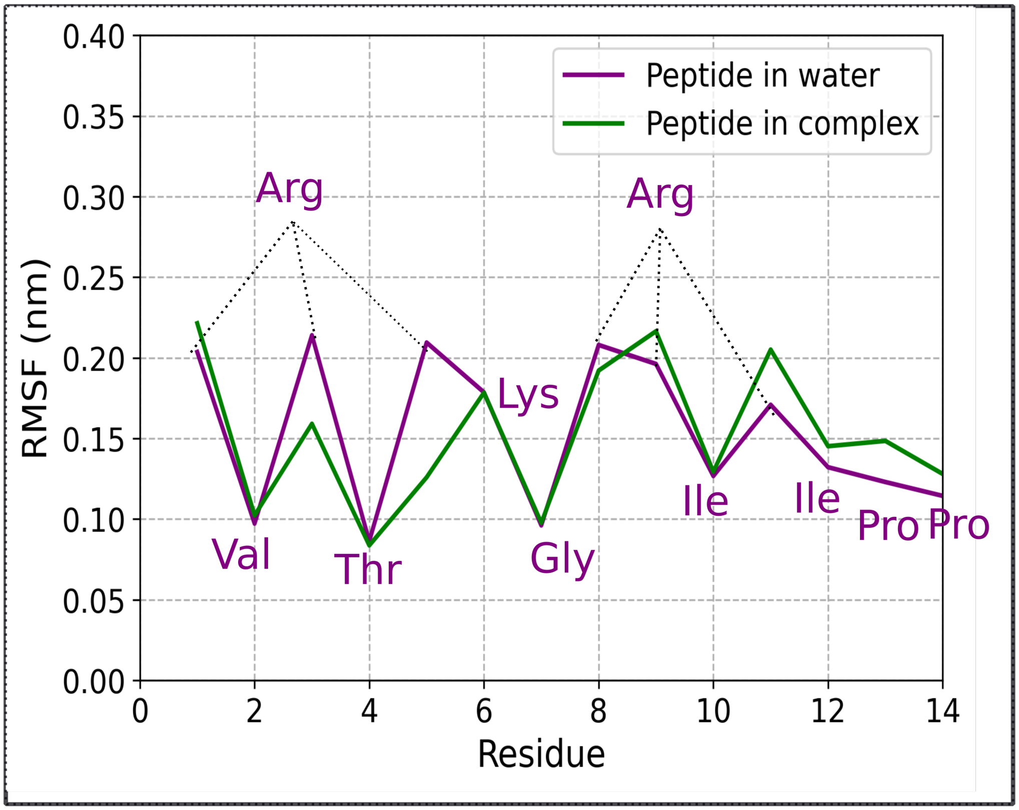


Figure S2: Root mean square fluctuation (RMSF) calculated using all the atoms of the Cyclic peptide.

**OPES colvar plot**

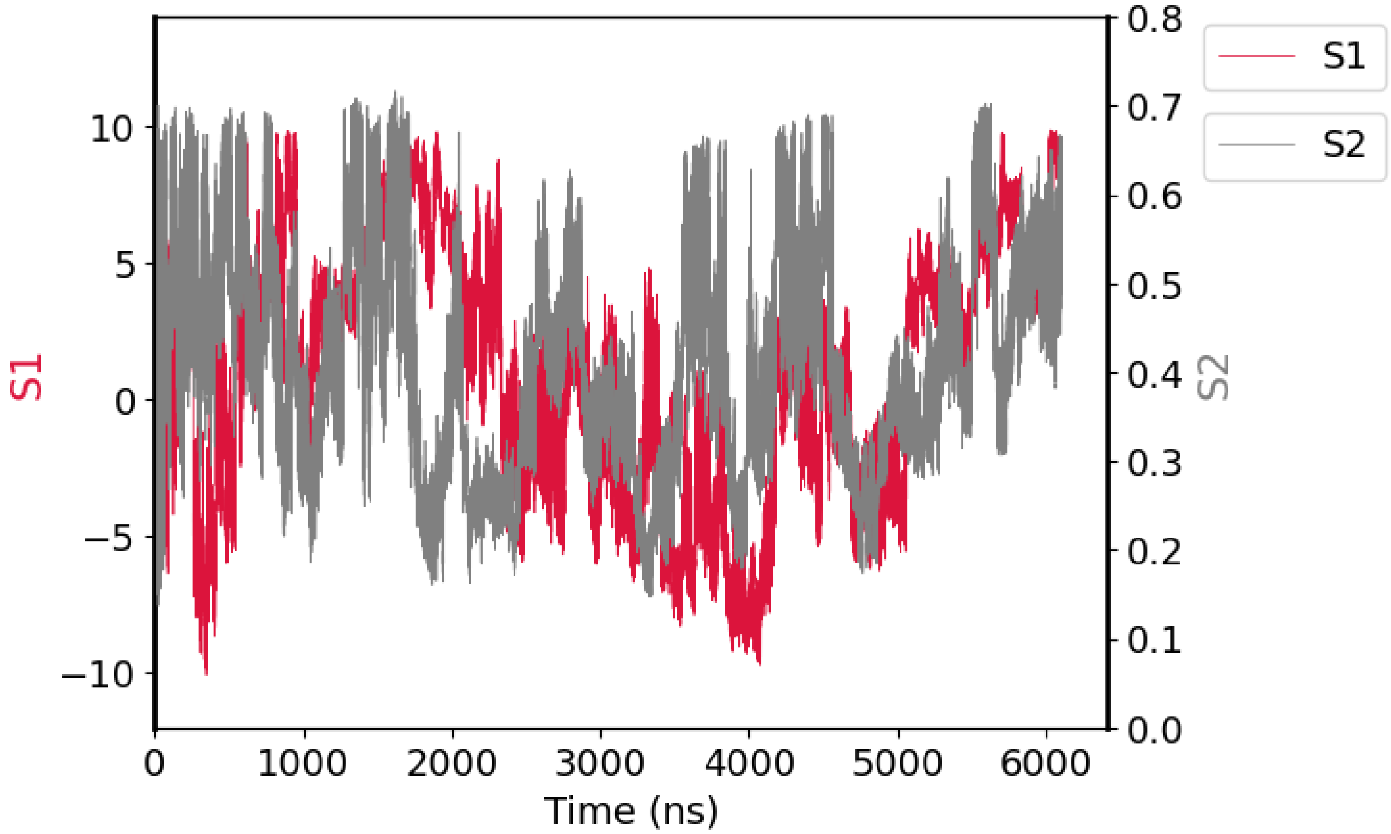


Figure S3: Plot of Deep-TDA and RMSD CV with simulation time. S1 corresponds to the deep-TDA CV. The bound and unbound states of the complex are represented by the two extremes of the Deep-TDA CV (y1-axis). Negative 10 corresponds to the bound state while positive 10 corresponds to the unbound state. Conversely, the y2-axis represents the RNA apical loop RMSD variations as the system investigates various metastable states.
